# Structural insights into Salinosporamide A mediated inhibition of the human 20S proteasome

**DOI:** 10.1101/2025.01.28.635221

**Authors:** Hagen Sülzen, Pavla Fajtova, Anthony J O’Donoghue, Evzen Boura, Jan Silhan

**Author notes:** Corresponding authors. Jan Silhan,; Evzen Boura.

## Abstract

The 20S proteasome, a critical component of the ubiquitin-proteasome system, plays a central role in regulating protein degradation in eukaryotic cells. Marizomib (MZB), a natural γ-lactam-β-lactone compound derived from *Salinispora tropica*, is a potent 20S proteasome covalent inhibitor with demonstrated anticancer properties. Its broad-spectrum inhibition of all three proteasome subunits and ability to cross the blood-brain barrier has made it a promising therapeutic candidate for glioblastoma. Here, we present the cryo-EM structure of the human 20S proteasome in complex with MZB at 2.55 Å resolution. This structure reveals the binding mode of MZB to all six catalytic subunits within the two β-rings of the 20S proteasome, providing a detailed molecular understanding of its irreversible inhibitory mechanism. These findings explain the therapeutic potential of MZB at the molecular level and highlight marine-derived natural products in targeting the proteasome for anticancer treatment.

## INTRODUCTION

Marizomib (MZB), also known as salinosporamide A, is a natural γ-lactam-β-lactone compound isolated from the marine bacterium *Salinispora tropica*.^1^ It is a non-peptidic 20S proteasome covalent inhibitor^1,2^ that has garnered significant attention for its potent anticancer properties and blood-brain barrier permeability.^3-5^ This unique capability makes marizomib particularly promising for the treatment of glioblastoma ^3^ and other central nervous system malignancies.

The proteasome is a critical component of the ubiquitin-proteasome system, which regulates protein degradation in eukaryotic cells.^6,7^ Structurally, the proteasome adopts a barrel-like architecture consisting of four heptameric rings, each formed by seven α or β subunits.^8-10^ The inner two rings harbor six catalytically active subunits (β1, β2, and β5) that possess N-terminal threonine residues critical for their enzymatic function.^11^ Despite the conserved architecture of these active sites, the substrate binding pockets preferentially cleave on the C-terminal side of hydrophobic amino acids (β5), positively charged residues (β2) or negatively charged residues (β1). These variations confer chymotrypsin-like, trypsin-like, and caspase-like substrate specificity to the proteasome, respectively.^12^

As proteasome activity is critical to numerous cellular processes, its dysfunction is associated with a variety of diseases.^13,14^ Cancer cells, characterized by rapid growth and genetic instability, are particularly dependent on proteasome activity to manage the large quantities of aberrant proteins they produce. Proteasome inhibitors like MZB disrupt this process, leading to the accumulation of misfolded proteins, cellular stress, and ultimately apoptosis. This mechanism of action is especially effective in cancer cells, which are more reliant on proteasome function than normal cells.

Marizomib irreversibly binds to the catalytic threonine of the 20S proteasome (**Figure 1**). As it lacks a peptide moiety, MZB does not bind in a substrate-like manner and therefore is capable of targeting all three subunits.^15-17^ MZB has highest affinity to the β5 subunit but readily binds to the β1 and β2 subunits at higher concentrations.^18,19^ This broad inhibition profile is likely to contribute to its potent anticancer activity, even in tumors resistant to other proteasome inhibitors. Furthermore, the unique β-lactone pharmacophore is highly specific for reacting with the catalytic threonine residues, as no other human hydrolases have been shown to be targeted by MZB. In addition to its anti-cancer effects, MZB is being investigated for applications in infectious diseases, such as malaria, by targeting the proteasome of *Plasmodium falciparum*.^20^

**Figure 1.**
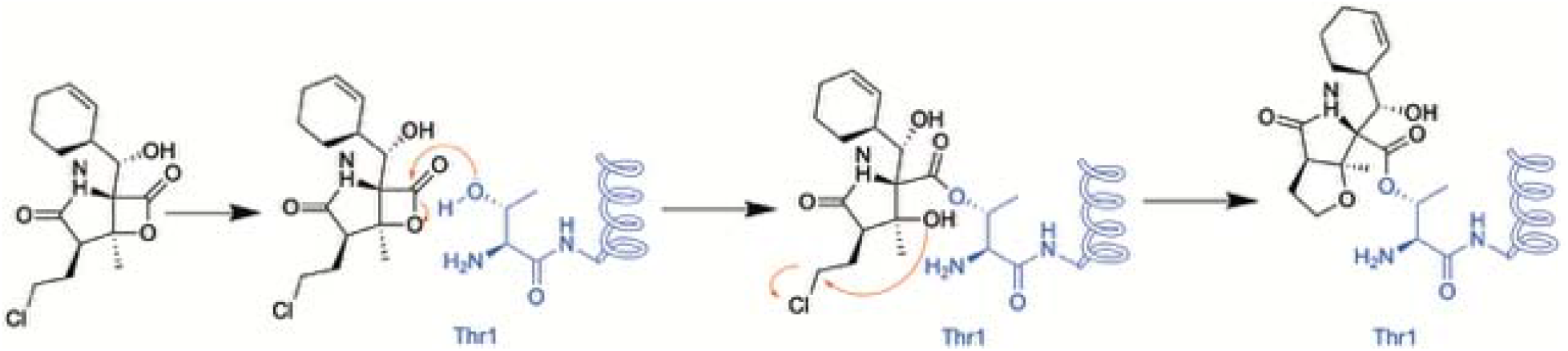
Schematic of the putative reaction mechanism between MZB and the N-terminal threonine of proteasome subunits.

Marizomib exemplifies the therapeutic potential of oceanic biodiversity in drug discovery. Beyond its ability to cross the blood-brain barrier, it also has enhanced stability and bioavailability compared to other peptide-like proteasome inhibitors. Currently in Phase III clinical trials, MZB has been evaluated in combination with standard glioblastoma treatments, such as temozolomide-based radiochemotherapy. Recent trial results revealed that there was no significant improvement in overall survival or progression-free survival in patients with newly diagnosed glioblastoma. In addition, patients receiving MZB experienced more documented adverse events when compared to those receiving the standard temozolomide-based radiochemotherapy.^3^ However, ongoing research continues to explore the use of MZB in combination therapies to enhance efficacy and address resistance mechanisms (ClinicalTrials.gov: NCT02330562).

The molecular details of how MZB binds to all three catalytic β-subunits have not been described. Therefore, in this study, we used single-particle cryo-EM to provide a detailed molecular explanation of the binding interactions with the human 20S proteasome.

## MATERIALS AND METHODS

Human constitutive proteasome (h20S, #E-360, R&D Systems), marizomib (MZB, MedChemExpress), fluorogenic substrates Suc-LLVY-amc, Z-VLR-amc and Z-LLE-amc were purchased from Cayman Chemical. Substrates were dissolved in DMSO at a concentration of 10□mM and stored at −20°C. MZB and h20S were aliquoted and stored at −80°C.

### Half-maximal inhibitory concentration (IC_50_) assays

The inhibition of individual h20S subunits was confirmed using subunit-specific fluorogenic substrates for the β1, β2, and β5 subunits, namely Z-LLE-amc, Z-VLR-amc, and Suc-LLVY-amc. Kinetic assays were performed in 384-well plates using 1 nM h20S preincubated with 100 nM PA28α human activator. The PA28α activator was expressed in *E*.*coli* using the pSumo vector, as previously described.^21^ The assay was conducted in a reaction buffer containing 50 mM HEPES (pH 7.5) and 1 mM DTT, with substrate concentrations of 80 μM Z-LLE-amc, 30 μM Z-VLR-amc, and 65 μM Ac-LLVY-amc, in a final volume of 8 μL per well. Inhibitors were dispensed into the plates using an Echo650 liquid handler (Beckman Coulter). Pre-steady-state kinetic measurements were carried out for the irreversible inhibitor, and IC_50_ values were calculated 60 to 90 minutes after initiating the reaction. Data analysis was performed using GraphPad Prism software. All assays were conducted in triplicate using 384-well black plates (Nunc) at 37 °C. Fluorescence measurements were recorded with a Synergy HTX Multi-Mode Microplate Reader (BioTek, Winooski, VT), using excitation and emission wavelengths of 360 nm and 460 nm, respectively.

### Preparation of cryo-EM grids and data acquisition

h20S (aprox. 0.6□μM) in a buffer consisting of 20 mM HEPES pH 7.5, 50 mM NaCl, 0.25 mM THP was incubated with 50 µM MZB at room temperature for 1 h before cooling the sample on ice. Four□μL of the proteasome-inhibitor complex (0.45 mg/mL) was applied to freshly glow discharged Quantifoil R2/1 300-mesh copper grids (EM Sciences, Prod. No. Q350CR1). Excess sample was removed by blotting with a FEI Vitrobot Mark IV (Thermo Fisher Scientific) (4□°C, 100% humidity, blot force -5) before plunge freezing the grids in liquid ethane.

The sample was imaged on a Titan Krios G3i microscope (Thermo Fisher Scientific) equipped with a Gatan K3 detector and operated at 300 kV. 4625 multi-frame movies (40 frames with a total dose of 60 e^-^/Å^2^) were recorded at a magnification of 105,000x, yielding a final pixel size of 0.8336 Å. Data were collected using the EPU v 3.0.0 data collection software (Thermo Fisher).

### Cryo-EM data processing

All image processing steps were performed in cryoSPARC (v4.3.0 and later).^22^ Movies were motion and CTF corrected using the Patch Motion and Patch CTF correction jobs respectively. Initial particle picking was performed using the Gaussian Blob Picker, resulting in a total of 1,949,389 particle picks. After several rounds of iterative 2D classification, 16,120 particles were used to generate an ab initio model without enforcing symmetry (C1).

Selected 2D classes were subsequently used as templates for reference-based particle picking, yielding 2,091,825 particle locations. Particles were extracted with a 400 px box and fourfold binning applied, resulting in a final pixel size of 3.32 Å/px. After iterative 2D classification, 355,436 selected particles were re-extracted with a 400 px box and no binning applied. The re-extracted particles and the ab-initio model generated previously were used to perform a round of non-uniform refinement with C2 symmetry enforced, before subsequently subjecting the aligned particles to a 3D classification job with 6 classes, resulting in four practically empty junk classes (≤ 32 particles) and two classes resembling the h20S proteasome. The best 3D class, comprised of 209,994 particles, was selected and subjected to homogenous refinement with C2 symmetry enforced, followed by reference-based motion correction. 257 particles were rejected due to their proximity to the micrograph edge, the remaining 209,737 particles were subjected to global and local CTF refinement before performing a final homogenous refinement with C2 symmetry, resulting in the final reconstruction with an FSC_0.143_ of 2.55Å (**Extended Data Table 1**). Finally, the handedness of the resulting map was flipped using the Volume Tool utility within cryoSPARC before sharpening the map using the EMReady software (v1.2)^23^. An overview of the cryo-EM data processing workflow is shown in **Extended Data Figure 1**.

### Modelling and Refinement

A previously determined cryo-EM structure of the human 20S proteasome (PDB: 7PG9)^24^ was used as a starting model and initially docked into the sharpened density map using ChimeraX (v1.7.1).^25^ Next, the model was iteratively refined using a combination of automated real-space refinement using Phenix real_space_refine,^26^ manual refinement in Coot 0.9.8.95 ^27,28^ and structure optimization using the ISOLDE^29^ package in ChimeraX until satisfactory validation metric and map-correlation had been achieved. Refinement restraints for MZB were generated using Jligand.^30^ Model validation was performed with MolProbity.^31^ Visualisation was performed in ChimeraX and PyMOL 3.0.1 (Schrödinger, LLC.)

## RESULTS

### Biochemical validation of the human 20S proteasome

The commercially obtained human 20S proteasome (h20S) was evaluated in biochemical assays for catalytic activity using fluorogenic reporter substrates that are each specific for either β1, β2 or β5 (**Figure 2**). After confirming the activity of all three catalytic subunits, we assessed whether these activities could be inhibited in the presence of MZB. Using a concentration range of 0 to 12.5 μM of MZB, we could demonstrate that the β5 subunits were inhibited with an IC_50_ of 18.5 nM while the IC_50_ for the β2 and β1 were 326.5 nM and 596.6 nM, respectively (**Figure 2**). Under these conditions all subunits were completely inhibited with 12.5 μM of MZB. These studies validate the quality of both the enzyme and inhibitor for structural studies.

**Figure 2.**
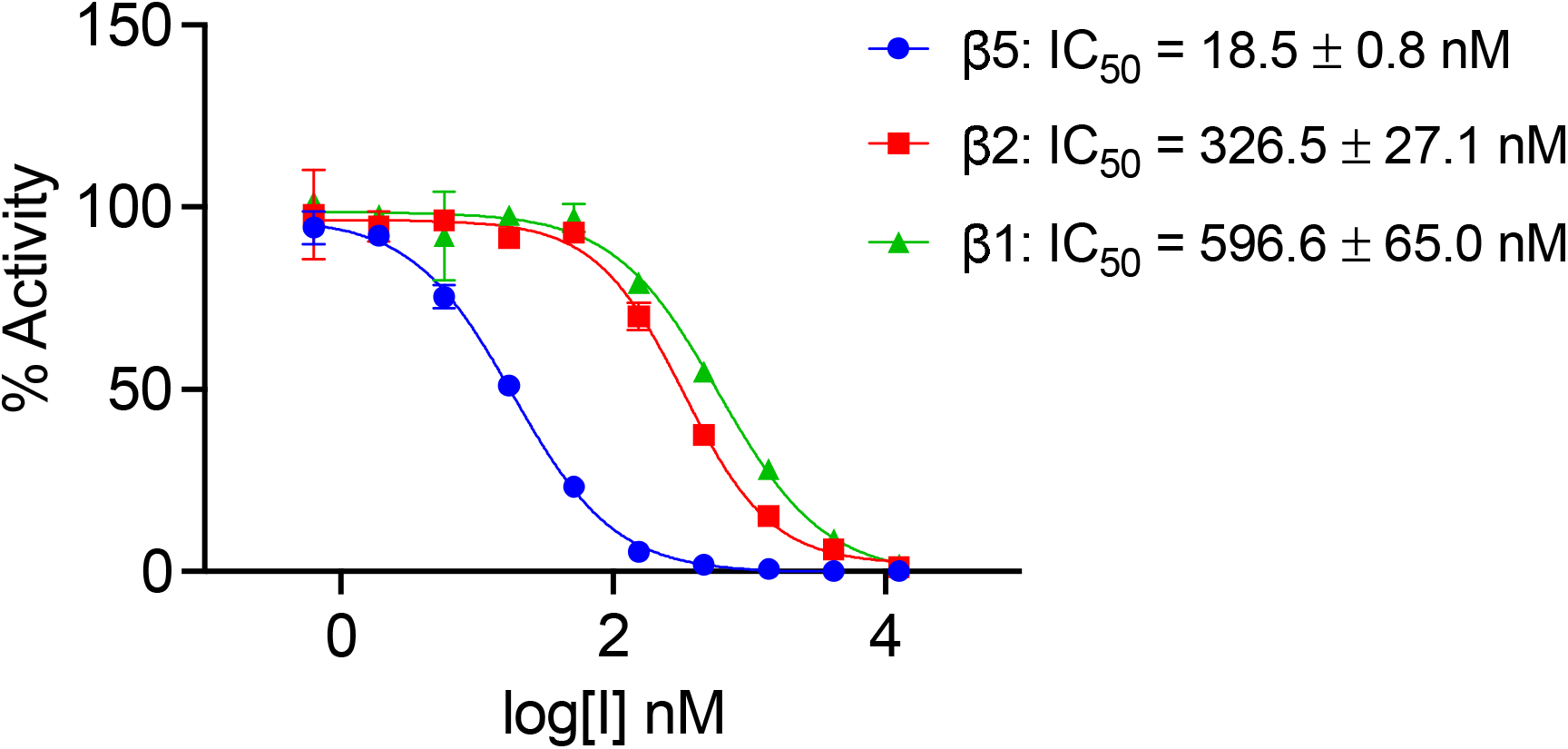
Half-maximal inhibitory concentration (IC_50_) of MZB for the proteolytic h20S subunits. IC_50_ curves for the inhibition of individual h20S proteasome subunits (β1, β2, and β5) determined using fluorogenic substrates Z-LLE-amc, Z-VLR-amc, and Suc-LLVY-amc. Assays were performed in 3 technical replicates and data are presented as mean values ± SD.

### Structural characterisation of the h20S proteasome

To gain structural insights into MZB mediated inhibition of the h20S, we employed single particle cryo-EM. The complex could be reconstructed to a reported FSC_0.143_ of 2.55 Å (**Figure 3A, Extended Data Figure 2**). While most particles observed in the micrographs appeared to present either side- or top and bottom views of the complex, continuous coverage of the Euler angle distribution allowed for a complete reconstruction (**Extended Data Figure 2B**). The local resolution of the reconstruction ranges predominantly between 2.53 and 5.7 Å (25^th^ to 75^th^ percentile), with lower resolutions observed primarily in regions corresponding to the solvent-facing surfaces of the α-subunits (**Extended Data Figure 2C**). The h20S proteasome could be modelled in its entirety except for flexible regions α1_Ala246-Asp249_, α2_Ala235-Ala237_, α3_Gln257-Lys262_, α4_Lys61_, α4_Glu241-Ser252_, α5_Val242-Ile246_, α6_Glu241-His266_, α7_Glu246-Met259_, β1_Pro203-Ala204_, β2_Val222-Ser233_, β4_Lys198-Ser201_, β5_Ser201-Pro203_, β6_Asp242_, β7_Gly262-Glu264_, covering 6256 of 6452 residues (96.96%) in total. An average map-to-model correlation of 0.92 (CC_side chain_, calculated with phenix.validation_cryoem^32^) indicates an excellent fit of the model to the experimental data (**Extended Data Figure 2D**).

**Figure 3.**
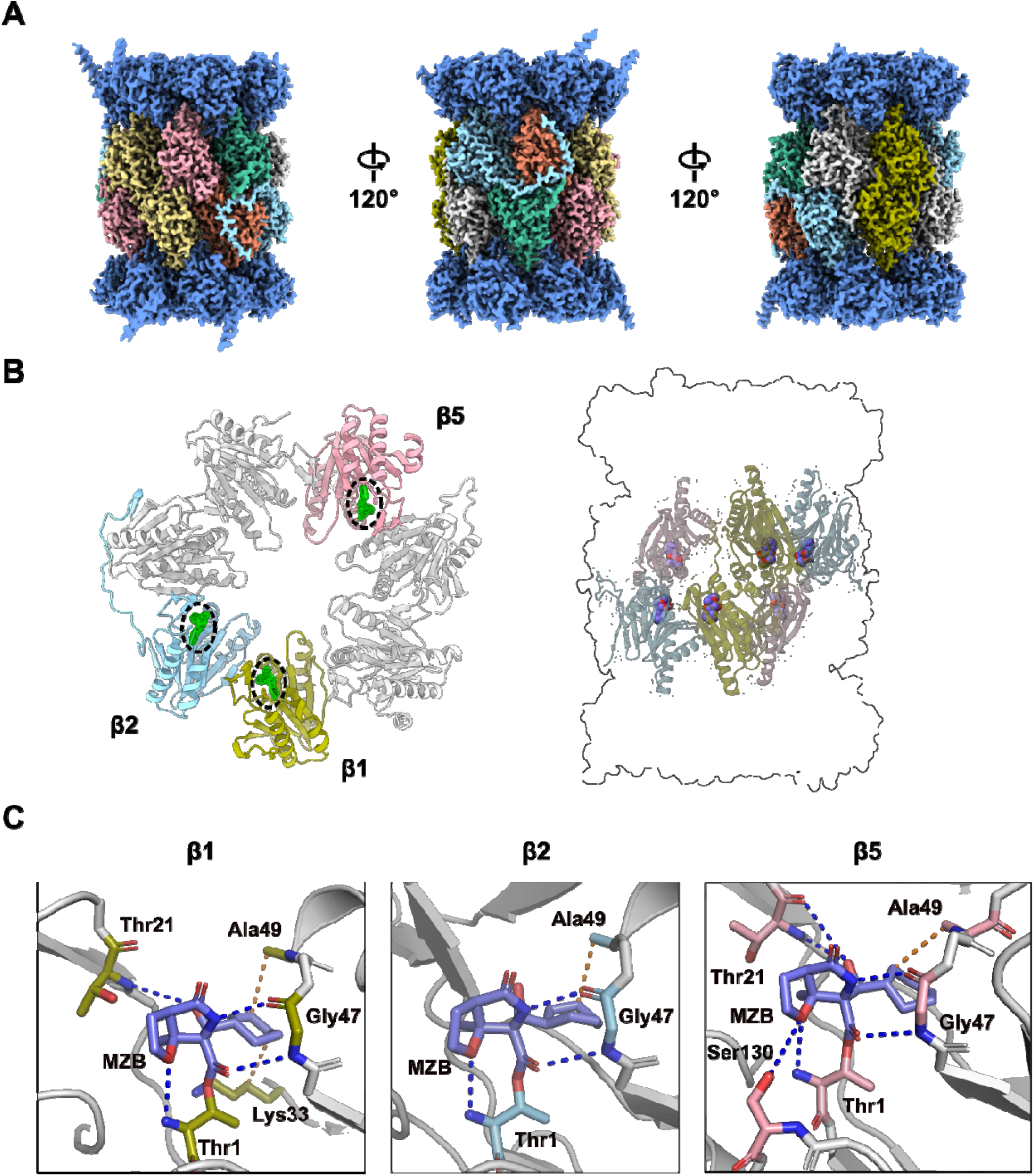
Cryo-EM structure of h20S in complex with MZB. **(A)** Final cryo-EM reconstruction after local sharpening using EMReady.^23^ Density corresponding to the α-subunits is coloured dark blue, proteolytic subunits β1, β2 and β5 are coloured in olive, cyan and pink, respectively. The remaining β subunits β3, β4, β6 and β7 are coloured in orange, yellow, mint and grey respectively. **(B)** Cartoon representation of one β subunit ring (left) and placement of the catalytic subunits relative to the overall proteasome architecture (right). The catalytic subunits follow the same colour scheme as in (A), the remaining subunits are coloured in light grey. The cryo-EM density for MZB is shown as green mesh (left) and the atomic model is depicted as spheres (right). **(C)** Close-up views of the three distinct active sites in β1, β2 and β5. MZB and interacting residues are displayed as coloured sticks. The colour scheme is consistent with (B). Side chains of non-interacting residues (white) have been removed for clarity. Hydrophobic interactions are displayed as orange dashed lines; hydrogen bonds are shown as blue dashed lines. Interaction analysis and initial visualisation were performed using PLIP.^34^

As expected, the atomic model obtained for the h20S proteasome in complex with MZB closely resembles the characteristic architecture commonly shared amongst 20S proteasomes and displays C2 symmetry, with the two sets of 14 subunits arranged in the conventional ring configuration of α1–α7, β1–β7 / β1–β7, α1–α7. Unsurprisingly, the h20S/MZB model could be aligned to the h20S structure used as a starting model (PDB: 7PG9) with an RMSD of 0.97 (27449 atoms), confirming an overall nearly identical fold.

### Binding of MZB to catalytic subunits β1, β2 and β5

The human 20S proteasome contains three proteolytically active subunits, β1, β2, and β5, within each β-subunit ring. Each active site features a conserved catalytic triad comprised of Thr1, Asp17 and Lys33, in addition to residues Ser129, Asp166 and Ser169, which are thought to provide structural integrity to the proteolytic centre and enhance catalysis.^33^ The well-defined density of the sharpened cryo-EM reconstruction allowed for unambiguous placement of the atomic model of the inhibitor (CC_Ligand_=0.93) (**Figure 3B**).

As expected, the carbonyl carbon atom derived from the beta-lactone ring of MZB is covalently bound to the oxygen from the hydroxyl side-chain group of the N-terminal catalytic Thr1 residue in all proteolytic subunits. Similarly, the orientation of the inhibitor is consistent across all three catalytic sites with the cyclohexenyl moiety occupying the so-called S1 pocket. While few hydrophobic interactions stabilize the ring structure in the S1 pocket, MZB is primarily coordinated via hydrogen bonding (**Figure 3C**). In all three active sites, residues Thr1 and Gly47 appear to coordinate the inhibitor via main chain interactions while for subunits β1 and β5 the cryo-EM reconstruction additionally places the peptide nitrogen (and carbonyl group) of residue Thr21 in coordination distance of the free carboxyl group in MZB (**Figure 3C**). Lastly, residue Ser130 in the catalytic site of the β5 subunit appears sufficiently close to coordinate the nitrogen of the γ-lactam ring of MZB via a sidechain interaction (**Figure 3C**). Intriguingly, the increased number of coordination sites for MZB in the active site of the β5 subunit correlates well with the higher potency of the inhibitor to this subunit (**Figure 2**). More detailed descriptions of the interactions between MZB and the active site pockets are shown in **Extended Data Figure 3**.

### Comparison of MZB binding to the active proteasome sites of human and T. vaginalis

MZB is a pan-proteasome inhibitor and has been suggested as a potential treatment for diseases caused by eukaryotic parasites. Recently, we determined the structure of the *Trichomonas vaginalis* 20S proteasome in complex with MZB.^19^ We compared how MZB binds these two distinct proteasomes, as these differences could be significant for future drug design.

While both, the human and *T. vaginalis* 20S proteasome (*Tv*20S), generally exhibit the commonly shared and characteristic architecture, structural alignment across all atoms reveals a root mean square deviation (RMSD) of 3.184 Å (**Figure 4A**), a surprising discrepancy in light of the rather high conservation of the individual subunits, ranging from 25-51% sequence identity.^19,35^ The largest differences between the atomic models are observed in the the α-subunit ring (**Figure 4A**). This deviation could possibly be attributed to the formation of filamentous structures by h20S upon vitrification on cryo-EM grids, where the α-subunit rings create the interface between individual particles (**Extended Data Figure 1**), potentially inducing conformational changes. In contrast, the *Tv*20S sample remained largely monodisperse.^19^ Alternatively, differences in the overall packing of the individual subunits could result in structural differences reflected by the inflated RMSD. Individual structural alignment of the proteolytic subunits results in RMSD values between 0.78 and 0.87 Å (**Figure 4A**), accentuating the conserved nature of the active sites.

**Figure 4.**
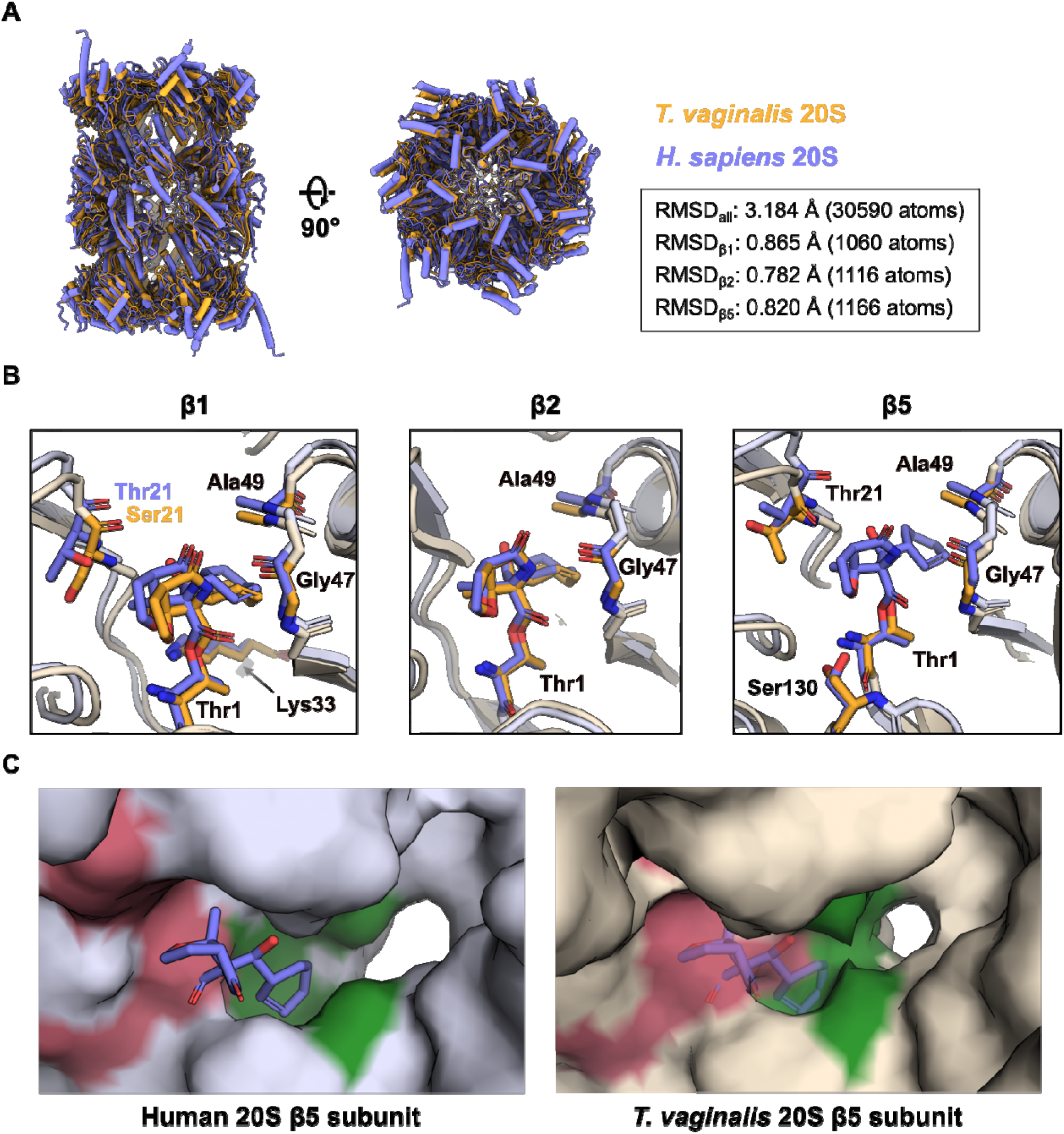
Comparison of h20S and *Tv*20S in complex with MZB. **(A)** Alignment and structural comparison of human (purple, PDB: 9HMN) and *T. vaginalis* (orange, PDB: 8OIX) 20S proteasomes bound to MZB. RMSD values for the alignment of the whole molecule and the individual catalytic subunits were obtained via PyMOL align with 5 cycles of outlier rejection. **(B)** Close-up views of the β1, β2 and β5 active sites. MZB and interacting residues are displayed as coloured sticks. The colour scheme is consistent with (A). Side chains of non-interacting residues (light purple/orange) have been removed for clarity. Labels for identical amino acids across both complexes are shown in black, deviating amino acids are shown in their respective colour. **(C)** Surface representation of the β5 subunit of human (left) and *T. vaginalis* (right) 20S proteasome. The catalytic site and S1 pocket are coloured pink and green, respectively. The atomic model for the *Tv*20S β5 subunit (PDB: 8OIX) does not include a model of MZB. Instead, MZB bound to the human protein has been aligned, superimposed and is shown as transparent sticks for representative purposes.

Indeed, structural comparison of the β1 and β2 subunits reveal nearly identical orientation and coordination of MZB (**Figure 4B**). While residue Thr21 of the h20 β1 subunit, predicted to form a hydrogen bond with MZB (**Figure 4C**), is replaced by Ser21 in the *T. vaginalis* equivalent, the peptide nitrogen is within bonding distance to the free carboxyl group of the inhibitor as well (3.7 Å, data not shown).

Intriguingly, our previous cryo-EM reconstruction of *Tv*20S had insufficient density to include MZB in the final model of the β5 subunit, despite the fact that the inhibitor attenuated proteolytic activity almost entirely *in vitro*.^19^ Alignment of the active site to the human β5 subunit once more reveals a nearly identical arrangement (**Figure 4B**), indicating structural differences elsewhere. As described earlier, in all three proteolytic subunits of h20S, the cyclohexenyl moiety of the inhibitor occupies the S1 pocket. When comparing the overall structural arrangement of the β5 subunits, it is apparent that access to the S1 cavity is much more restricted in *T. vaginalis* (**Figure 4C**). It is conceivable that, as a result of this steric impairment, a reduced number of coordination sites are available, granting the covalently bound MZB increased flexibility and thereby smearing its cryo-EM density.

## DISCUSSION

Proteasomal degradation of ubiquitinated proteins is a key cellular process that controls protein turn over, and thus plays a crucial role in numerous cellular functions. These include development and differentiation, DNA repair, cell cycle progression, immune responses, apoptosis, and stress adaptation. The ability to selectively degrade specific targets is crucial to maintains cellular homeostasis and to dynamically regulate essential biological pathways.^36^ Proteasome inhibition disrupts these tightly regulated processes by preventing protein degradation, which induces cellular stress and, eventually, leads to apoptosis. This mechanism is particularly effective in selectively killing rapidly dividing cancer cells such as myeloma cells. These cells rely heavily on protein integrity to sustain their high production of gamma globulin.^37^ Consequently, FDA-approved proteasome inhibitors, such as bortezomib, carfilzomib, and ixazomib are used to treat multiple myeloma and other cancers.^38,39^ New and promising drugs, such as marizomib (MZB), oprozomib, and delanzomib, are currently undergoing testing, either as standalone treatments or in combination with other therapeutics.^40^

Structural information is crucial for guiding the development of new proteasome inhibitors with enhanced efficacy and specificity. The knowledge of MZB’s interactions with the human proteasome can help design novel therapeutics for various cancers. In this study, we investigated the binding of MZB to the human 20S proteasome using cryo-EM, revealing its molecular architecture and specific interactions. High-resolution cryo-EM maps confirmed that MZB irreversibly binds to all three catalytic subunits of the proteasome: chymotrypsin-like (β5), trypsin-like (β2), and caspase-like (β1). This pan-inhibitory activity distinguishes MZB from other proteasome inhibitors, such as bortezomib and carfilzomib, which primarily target the β5 site and may lead to compensatory activation of other sites, potentially contributing to resistance.^39^ Our findings provide structural insights at near-atomic resolution, explaining the broad inhibitory action of MZB. This knowledge can serve as a foundation for optimizing MZB’s application or redesigning it to achieve more targeted therapeutic effects.

Furthermore, using a combination of proteasome inhibitors with other drugs may pave the way for safer and more specific cancer therapies. Preclinical studies have demonstrated promising results for marizomib (MZB) in triple-negative breast cancer (TNBC), where it effectively inhibits primary tumor growth and metastasis to the lungs and brain, potentially addressing the limitations of existing therapies like bortezomib and carfilzomib.^41^ In cervical cancer, combining MZB with cisplatin shows synergistic effects both in vitro and in vivo, underscoring the need for further investigation into optimal dosing and scheduling.^42^ Additionally, the potential of MZB combinations with other anti-cancer agents, such as glycolysis inhibitors, is particularly compelling, given the observed upregulation of glycolysis pathway proteins following MZB treatment in TNBC cells.^41^ However, further research is required to evaluate the long-term efficacy of MZB, identify potential resistance mechanisms, and refine combination strategies with other therapeutics. Understanding how MZB impacts cell cycle progression, apoptosis, and protein degradation pathways will be critical, particularly for designing synergistic approaches. Moreover, addressing whether prolonged MZB exposure induces acquired resistance, as observed with previous proteasome inhibitors, will be crucial for maximizing its therapeutic potential.^39,43^ These efforts will be instrumental in optimizing MZB for the treatment of cancer and, potentially, to treat parasitic infections.^21,44^

Beyond cancer treatments, marizomib (MZB) shows potential for combating parasitic infections. Our comparative analysis of MZB binding to the human 20S proteasome and the eukaryotic parasite *Trichomonas vaginalis* (*Tv*20S) proteasome reveals a promising pathway for developing targeted treatments against eukaryotic parasites. Structural insights into the *Tv*20S proteasome suggest that MZB could be chemically modified to selectively target the β5 subunit of the *Tv*20S proteasome. Such modifications could result in a drug highly specific to *T. vaginalis*, offering a novel therapeutic option for diseases caused by this parasite while minimizing off-target effects on human cells.

In conclusion, elucidating MZB’s mechanism of action represents a critical step toward designing more effective therapeutics. MZB could be improved to more effectively treat cancer and, moreover, could be transformed into a drug to effectively treat eukaryotic parasites such as *T. vaginalis*.

## Data availability

The atomic model of the presented complex of human 20S proteasome with small molecule inhibitor MZB has been deposited in the Protein Data Bank under accession code 9HMN. The associated cryo-EM density maps, half-maps and masks have been deposited in the Electron Microscopy Data Bank under accession code EMD -52296. The starting model used for the modelling of the complex is deposited in the Protein Data Bank under accession code 7PG9.

## Acknowledgements

We thank Tomas Kouba and Anatolij Filimonenko for assisting in the collection of the cryo-EM data. This research was funded by the project the National Institute Virology and Bacteriology (Programme EXCELES, Project No. LX22NPO5103) -Funded by the European Union - Next Generation EU. We also acknowledge the CF BIC of CIISB, Instruct-CZ Centre, supported by MEYS CR (LM2023042) and European Regional Development Fund-Project "Innovation of Czech Infrastructure for Integrative Structural Biology” (No. CZ.02.01.01/00/23_015/0008175). P.F. received funding from the European Union’s Horizon 2020 research and innovation program under the Marie Skłodowska-Curie grant agreement No [846688]. The research was also supported by NIH awards R01AI158612, R21AI146387, R21AI133393 and R21AI171824 to A.J.O. We thank Dr. Jehad Almaliti for his explanation of the mechanism of action of MZB.

## Author contributions

J.S. and E.B. conceived the project. J.S. and H.S. collected and processed cryo-EM data, P.F. performed enzyme experiments. H.S., P.F, A.J.O., J.S., and E.B. wrote, revised and edited the manuscript. P.F, A.J.O., J.S., and E.B obtained funding.

## REFERENCES

1 Feling, R. H. et al. Salinosporamide A: a highly cytotoxic proteasome inhibitor from a novel microbial source, a marine bacterium of the new genus salinospora. Angew Chem Int Ed Engl 42, 355–357 (2003). 10.1002/anie.200390115

2 Mincer, T. J., Jensen, P. R., Kauffman, C. A. & Fenical, W. Widespread and persistent populations of a major new marine actinomycete taxon in ocean sediments. Appl Environ Microbiol 68, 5005–5011 (2002). 10.1128/AEM.68.10.5005-5011.2002

3 Roth, P. et al. Marizomib for patients with newly diagnosed glioblastoma: A randomized phase 3 trial. Neuro Oncol 26, 1670–1682 (2024). 10.1093/neuonc/noae053

4 Di, K. et al. Marizomib activity as a single agent in malignant gliomas: ability to cross the blood-brain barrier. Neuro Oncol 18, 840–848 (2016). 10.1093/neuonc/nov299

5 Manton, C. A. et al. Induction of cell death by the novel proteasome inhibitor marizomib in glioblastoma in vitro and in vivo. Sci Rep 6, 18953 (2016). 10.1038/srep18953

6 Manasanch, E. E. et al. The proteasome: mechanisms of biology and markers of activity and response to treatment in multiple myeloma. Leuk Lymphoma 55, 1707–1714 (2014). 10.3109/10428194.2013.828351

7 Eisenberg-Lerner, A. et al. Golgi organization is regulated by proteasomal degradation. Nat Commun 11, 409 (2020). 10.1038/s41467-019-14038-9

8 Tanaka, K. The proteasome: overview of structure and functions. Proc Jpn Acad Ser B Phys Biol Sci 85, 12–36 (2009). 10.2183/pjab.85.12

9 Coux, O., Tanaka, K. & Goldberg, A. L. Structure and functions of the 20S and 26S proteasomes. Annu Rev Biochem 65, 801–847 (1996). 10.1146/annurev.bi.65.070196.004101

10 Lupas, A., Zwickl, P., Wenzel, T., Seemuller, E. & Baumeister, W. Structure and function of the 20S proteasome and of its regulatory complexes. Cold Spring Harb Symp Quant Biol 60, 515–524 (1995). 10.1101/sqb.1995.060.01.055

11 Lowe, J. et al. Crystal structure of the 20S proteasome from the archaeon T. acidophilum at 3.4 A resolution. Science 268, 533–539 (1995). 10.1126/science.7725097

12 Orlowski, M., Cardozo, C. & Michaud, C. Evidence for the presence of five distinct proteolytic components in the pituitary multicatalytic proteinase complex. Properties of two components cleaving bonds on the carboxyl side of branched chain and small neutral amino acids. Biochemistry 32, 1563–1572 (1993). 10.1021/bi00057a022

13 Mishra, R., Upadhyay, A., Prajapati, V. K. & Mishra, A. Proteasome-mediated proteostasis: Novel medicinal and pharmacological strategies for diseases. Med Res Rev 38, 1916–1973 (2018). 10.1002/med.21502

14 Schmidt, M. & Finley, D. Regulation of proteasome activity in health and disease. Biochim Biophys Acta 1843, 13–25 (2014). 10.1016/j.bbamcr.2013.08.012

15 Fenteany, G. et al. Inhibition of proteasome activities and subunit-specific amino-terminal threonine modification by lactacystin. Science 268, 726–731 (1995). 10.1126/science.7732382

16 Groll, M. & Potts, B. C. Proteasome structure, function, and lessons learned from beta-lactone inhibitors. Curr Top Med Chem 11, 2850–2878 (2011). 10.2174/156802611798281320

17 Macherla, V. R. et al. Structure-activity relationship studies of salinosporamide A (NPI-0052), a novel marine derived proteasome inhibitor. J Med Chem 48, 3684–3687 (2005). 10.1021/jm048995+

18 Groll, M., Huber, R. & Potts, B. C. Crystal structures of Salinosporamide A (NPI-0052) and B (NPI-0047) in complex with the 20S proteasome reveal important consequences of beta-lactone ring opening and a mechanism for irreversible binding. J Am Chem Soc 128, 5136–5141 (2006). 10.1021/ja058320b

19 Silhan, J. et al. Structural elucidation of recombinant Trichomonas vaginalis 20S proteasome bound to covalent inhibitors. Nature communications 15, 8621 (2024). 10.1038/s41467-024-53022-w

20 Prudhomme, J. et al. Marine actinomycetes: a new source of compounds against the human malaria parasite. PLoS One 3, e2335 (2008). 10.1371/journal.pone.0002335

21 Robbertse, L. et al. Evaluating Antimalarial Proteasome Inhibitors for Efficacy in Babesia Blood Stage Cultures. ACS Omega 9, 44989–44999 (2024). 10.1021/acsomega.4c04564

22 Punjani, A., Rubinstein, J. L., Fleet, D. J. & Brubaker, M. A. cryoSPARC: algorithms for rapid unsupervised cryo-EM structure determination. Nature methods 14, 290–296 (2017). 10.1038/nmeth.4169

23 He, J., Li, T. & Huang, S. Y. Improvement of cryo-EM maps by simultaneous local and non-local deep learning. Nature communications 14, 3217 (2023). 10.1038/s41467-023-39031-1

24 Sahu, I. et al. The 20S as a stand-alone proteasome in cells can degrade the ubiquitin tag. Nat Commun 12, 6173 (2021). 10.1038/s41467-021-26427-0

25 Pettersen, E. F. et al. UCSF ChimeraX: Structure visualization for researchers, educators, and developers. Protein science : a publication of the Protein Society 30, 70–82 (2021). 10.1002/pro.3943

26 Afonine, P. V. et al. Real-space refinement in PHENIX for cryo-EM and crystallography. Acta Crystallogr D Struct Biol 74, 531–544 (2018). 10.1107/S2059798318006551

27 Liebschner, D. et al. Macromolecular structure determination using X-rays, neutrons and electrons: recent developments in Phenix. Acta Crystallogr D Struct Biol 75, 861–877 (2019). 10.1107/S2059798319011471

28 Emsley, P., Lohkamp, B., Scott, W. G. & Cowtan, K. Features and development of Coot. Acta crystallographica. Section D, Biological crystallography 66, 486–501 (2010). 10.1107/S0907444910007493

29 Croll, T. I. ISOLDE: a physically realistic environment for model building into low-resolution electron-density maps. Acta Crystallogr D Struct Biol 74, 519–530 (2018). 10.1107/S2059798318002425

30 Lebedev, A. A. et al. JLigand: a graphical tool for the CCP4 template-restraint library. Acta Crystallogr D Biol Crystallogr 68, 431–440 (2012). 10.1107/S090744491200251X

31 Williams, C. J. et al. MolProbity: More and better reference data for improved all-atom structure validation. Protein Sci 27, 293–315 (2018). 10.1002/pro.3330

32 Afonine, P. V. et al. New tools for the analysis and validation of cryo-EM maps and atomic models. Acta Crystallogr D Struct Biol 74, 814–840 (2018). 10.1107/S2059798318009324

33 Borissenko, L. & Groll, M. 20S proteasome and its inhibitors: crystallographic knowledge for drug development. Chem Rev 107, 687–717 (2007). 10.1021/cr0502504

34 Adasme, M. F. et al. PLIP 2021: expanding the scope of the protein-ligand interaction profiler to DNA and RNA. Nucleic Acids Res 49, W530–W534 (2021). 10.1093/nar/gkab294

35 Fajtova, P. et al. Distinct substrate specificities of the three catalytic subunits of the Trichomonas vaginalis proteasome. Protein science : a publication of the Protein Society 33, e5225 (2024). 10.1002/pro.5225

36 Ciechanover, A. The Ubiquitin-Proteasome Proteolytic Pathway. Cell 79, 13–21 (1994). 10.1016/0092-8674(94)90396-4

37 Chauhan, D. et al. A novel orally active proteasome inhibitor induces apoptosis in multiple myeloma cells with mechanisms distinct from Bortezomib. Cancer cell 8, 407–419 (2005). 10.1016/j.ccr.2005.10.013

38 Nunes, A. T. & Annunziata, C. M. Proteasome inhibitors: structure and function. Semin Oncol 44, 377–380 (2017). 10.1053/j.seminoncol.2018.01.004

39 Park, J. E., Miller, Z., Jun, Y., Lee, W. & Kim, K. B. Next-generation proteasome inhibitors for cancer therapy. Translational Research 198, 1–16 (2018). 10.1016/j.trsl.2018.03.002

40 Kegyes, D. et al. Proteasome inhibition in combination with immunotherapies: State-of-the-Art in multiple myeloma. Blood Rev 61 (2023). https://doi.org:ARTN10110010.1016/j.blre.2023.101100

41 Raninga, P. et al. Marizomib suppresses triple-negative breast cancer via proteasome and oxidative phosphorylation inhibition. Theranostics 10, 5259–5275 (2020). 10.7150/thno.42705

42 Zhang, Z. et al. Combined treatment of marizomib and cisplatin modulates cervical cancer growth and invasion and enhances antitumor potential in vitro and in vivo. Front Oncol 12, 974573 (2022). 10.3389/fonc.2022.974573

43 Kaplan, G. S., Torcun, C. C., Grune, T., Ozer, N. K. & Karademir, B. Proteasome inhibitors in cancer therapy: Treatment regimen and peripheral neuropathy as a side effect. Free Radic Biol Med 103, 1–13 (2017). 10.1016/j.freeradbiomed.2016.12.007

44 Eadsforth, T. C. et al. Pharmacological and structural understanding of the Trypanosoma cruzi proteasome provides key insights for developing site-specific inhibitors. The Journal of biological chemistry 301, 108049 (2024). 10.1016/j.jbc.2024.108049

