## Supplementary Figures and Tables for "Structural insights into Salinosporamide A mediated inhibition of the human 20S proteasome"

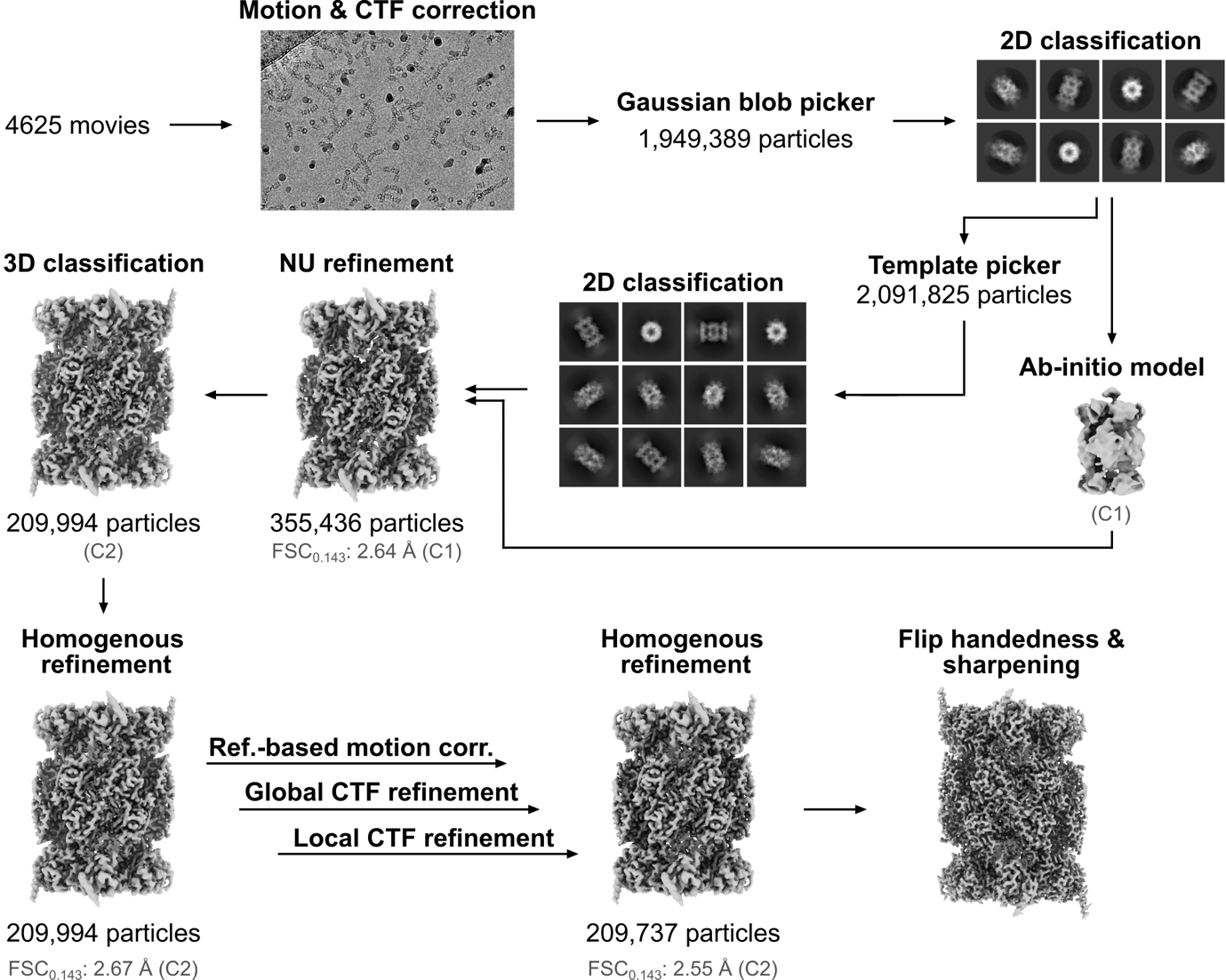

**Supplementary Figure 1. Workflow of cryo-EM data processing.** Visual representation of the image processing workflow used to reconstruct the h20S proteasome in complex with marizomib. A representative micrograph and selected 2D classes are shown in addition to the ab-initio model used and the density maps that were obtained for each reconstruction step. Where applicable, the number of particles, the gold-standard FSC estimate of the reconstruction resolution and the symmetry applied in the processing step (shown in brackets) are indicated. A total of 209,737 particles were used for the final reconstruction. All image processing steps were performed using cryoSPARC.^1^ The final sharpening step was performed using EMReady.^2^

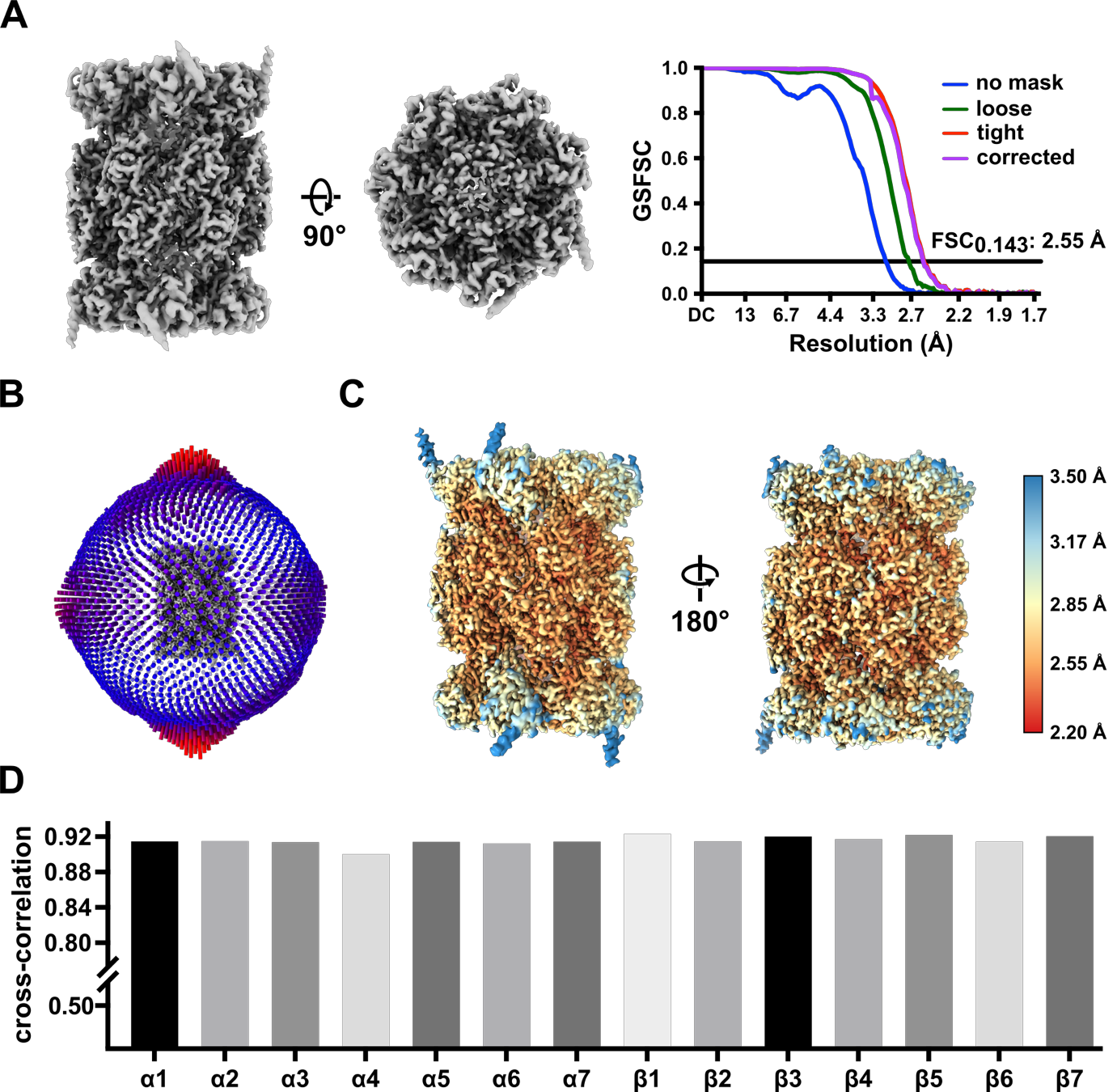

**Supplementary Figure 2. Cryo-EM single particle analysis of h20S in complex with MZB. (A)** Final (unsharpened) cryo-EM reconstruction (left) used to calculate the gold-standard Fourier-shell correlation curves (right). **(B)** Angular distribution plot of the final reconstruction. The height of the bars corresponds to the relative abundance of particle views contributing to the final reconstruction. The apparent symmetric distribution of Euler angles is the result of enforcing C2 symmetry during the reconstruction. **(C)** Colour-graded representation of the local resolution (in Å) of the final reconstruction, visualised on the EMReady sharpened map with corrected handedness. The range of the colour gradient was chosen to achieve the best visual representation and does not cover all spatial frequencies present in the reconstruction. **(D)** Average map-to-model cross-correlation (CC_side chain_) for h20S subunits. Since the atomic models plotted here have been fully refined prior to symmetry expansion, the symmetry-related copies are not included here.

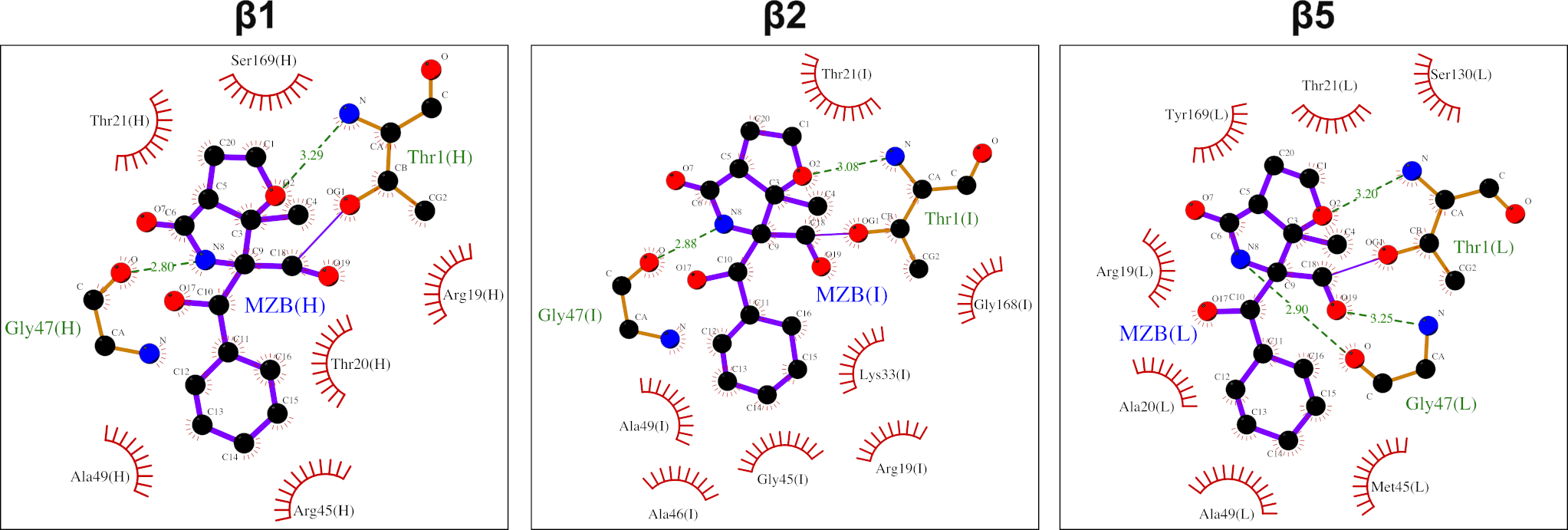

**Supplementary Figure 3. 2D representation of the interactions between small molecule inhibitor MZB and the h20S active sites.** Hydrogen bonds are displayed as green dashed lines, non-ligand residues involved in hydrophobic contacts are displayed as red semicircular arcs. Plots were generated using LigPlot.^3^

**Supplementary Table 1: Cryo-EM data collection, refinement and validation statistics.**

|  | h20S-MZB  (EMDB-52296)  (PDB- 9HMN) |
| --- | --- |
| **Data collection and processing** |  |
| Microscope | Titan Krios |
| Detector | Gatan K3 |
| Magnification (nominal) | 105.000x |
| Voltage (kV) | 300 |
| Spherical aberration | 2.7 mm |
| Total electron dose (e–/Å^2^) | 60 |
| Defocus range (μm) | -2.5 to -1.0 |
| Pixel size (Å) | 0.8336 |
| Stage tilt | 0° |
| Number of Micrographs | 4625 |
| Final particle images (no.) | 209,737 |
| Map resolution (Å)  [FSC threshold] | 2.55 [FSC_0.143_] |
| **Refinement** |  |
| Initial model used (PDB code) | 7PG9 |
| Symmetry during reconstruction | C2 |
| RMSD  Bond lengths (Å)  Bond angles (°) | 0.007  1.082 |
| Validation  MolProbity score  Clashscore, all-atom  Rotamer outliers | 1.27  5.1  0.88% |
| Ramachandran plot  Favoured  Allowed  Outliers | 98.05%  1.76%  0.19% |
| **Model vs. Data** |  |
| Ligands (no.) | 6 (MZB) |
| CC (mask/box/ligand) | 0.91/ 0.94 / 0.93 |
| Resolution estimates (Å)  FSC (0.143/0.5) | 2.55 / 2.8 |

**Supplementary references**

1	Punjani, A., Rubinstein, J. L., Fleet, D. J. & Brubaker, M. A. cryoSPARC: algorithms for rapid unsupervised cryo-EM structure determination. *Nature methods* **14**, 290-296 (2017). https://doi.org:10.1038/nmeth.4169

2	He, J., Li, T. & Huang, S. Y. Improvement of cryo-EM maps by simultaneous local and non-local deep learning. *Nature communications* **14**, 3217 (2023). https://doi.org:10.1038/s41467-023-39031-1

3	Laskowski, R. A. & Swindells, M. B. LigPlot+: multiple ligand-protein interaction diagrams for drug discovery. *Journal of chemical information and modeling* **51**, 2778-2786 (2011). https://doi.org:10.1021/ci200227u
